# To 200,000 m/z and beyond: native electron capture charge reduction mass spectrometry deconvolves heterogeneous signals in large biopharmaceutical analytes

**DOI:** 10.1101/2024.02.19.581059

**Authors:** Kyle I. P. Le Huray, Tobias P. Wörner, Tiago Moreira, Maria Reinhardt-Szyba, Paul W. A. Devine, Nicholas J. Bond, Kyle L. Fort, Alexander A. Makarov, Frank Sobott

**Author notes:** These authors contributed equally (Kyle I. P. Le Huray, Tobias P. Wörner).

## Abstract

Great progress has been made in the detection of large biomolecular analytes by native mass spectrometry, however characterizing highly heterogeneous samples remains challenging due to the presence of many overlapping signals from complex ion distributions. Electron-capture charge reduction (ECCR), in which a protein cation captures free electrons without apparent dissociation, can dealign overlapping signals by shifting the ions to lower charge states. The concomitant shift to higher *m/z* also facilitates the exploration of instrument upper *m/z* limits if large complexes are used. Here we perform native ECCR on the bacterial chaperonin GroEL and megadalton scale adeno-associated virus (AAV) capsid assemblies on the Q Exactive UHMR mass spectrometer. Charge reduction of AAV8 capsids by up to 90% pushes signals well above 100,000 *m/z* and enables charge state resolution and mean mass determination of these highly heterogeneous samples, even for capsids loaded with genetic cargo. With minor instrument modifications the UHMR can detect charge reduced ion signals beyond 200,000 *m/z*. This work demonstrates the utility of ECCR for deconvolving heterogeneous signals in native mass spectrometry and presents the highest *m/z* signals ever recorded on an Orbitrap instrument, opening up the use of Orbitrap native mass spectrometry for heavier analytes than ever before.

## Introduction

Native mass spectrometry involves the ionization and solution-to-gas phase transfer of biological macromolecules and complexes such as proteins and nucleic acids, while preserving a near-native structure and maintaining non-covalent interactions (*1–4*). Biological assemblies studied in this way range from small protein-ligand complexes to diverse and heterogeneous ribosomes, intact viruses, DNA nanostructures and protein-lipid supercomplexes released directly from membranes (*5–10*). The number of charges acquired by a biomolecular analyte during nano-electrospray ionization (nano-ESI) depends primarily on its solution-phase solvent accessible surface area and the chemical composition of the solution (*11, 12*). Manipulation of the charge state distribution, either to reduce (charge reduction) or to increase (supercharging) the number of charges on the analyte can be desirable in multiple use-cases. In cases where the analyte natively charges by nano-ESI at a mass-to-charge ratio (*m/z*) range beyond the capabilities of an instrument, charge manipulation can be used to move the m/z of the analyte into the detectable range (*13*). Electric fields are used to control ions in the mass spectrometer, and in combination with a background collision gas, ions can be accelerated to cause collisional energy transfer, for example to achieve fragmentation, dissociation or desolvation; the extent of acceleration and therefore activation at a given voltage is related to the ion’s charge, and charge manipulation can therefore help to control the extent of activation. Supercharging has for example been shown to assist in desolvation for some membrane proteins, for which greater activation is needed to eject the protein from detergent micelles; however, in other cases, charge reduction is useful to prevent unfolding of the protein due to over-activation or internal Coulomb repulsion, and is also of fundamental interest for understanding protein structure in the gas phase (*12, 14–18*).

A major use-case for charge reduction in native mass spectrometry is to attain separation of overlapping peaks resulting from the heterogeneity present in many biological macromolecules. The ideal native mass spectrum yields a fully resolved distribution of signals for each species over different charge states; assignment of ion charge from the distribution enables mass calculation, and ideally resolves mass heterogeneity resulting from, for example, ligand binding (Fig. 1A). As long as neighbouring charge states do not overlap with each other and the ions are sufficiently desolvated, a high resolving power of the mass analyser should enable the resolution of even the smallest mass differences, up to the isotopic fine structure within the analytes (*19*). However, as analytes increase in size and complexity, the mass differences between individual species can become difficult to resolve where the ions of neighbouring charge state distributions merge into each other, and ions of different charge states coincide exactly at the same *m/z* position ((*m*/*z*)_*ion*;1_ = (*m*/*z*)_*ion*;2_ = (*m*/*z*)_*shared*_), or at close enough *m/z* position to cause peak overlap unresolvable with the available resolving power of the mass analyzer. This is most often the case in samples where the analyte can harbour several modifications which increase the mass, so that it merges into the adjacent charge state and (equation 1) is fulfilled:

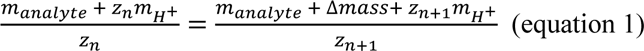

Where *m_analyte_* is the mass of the analyte of charge *z*_*n*_, which will have exact peak convolution (overlap) with the *z*_*n*+1_ charge state of an ion with additional mass, Δ*mass*. Rearranging this we can calculate at which *Δmass* complete peak overlap would occur, causing exact peak overlap which is unresolvable even at near-infinite resolving power:

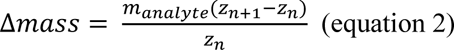

**Figure 1.**
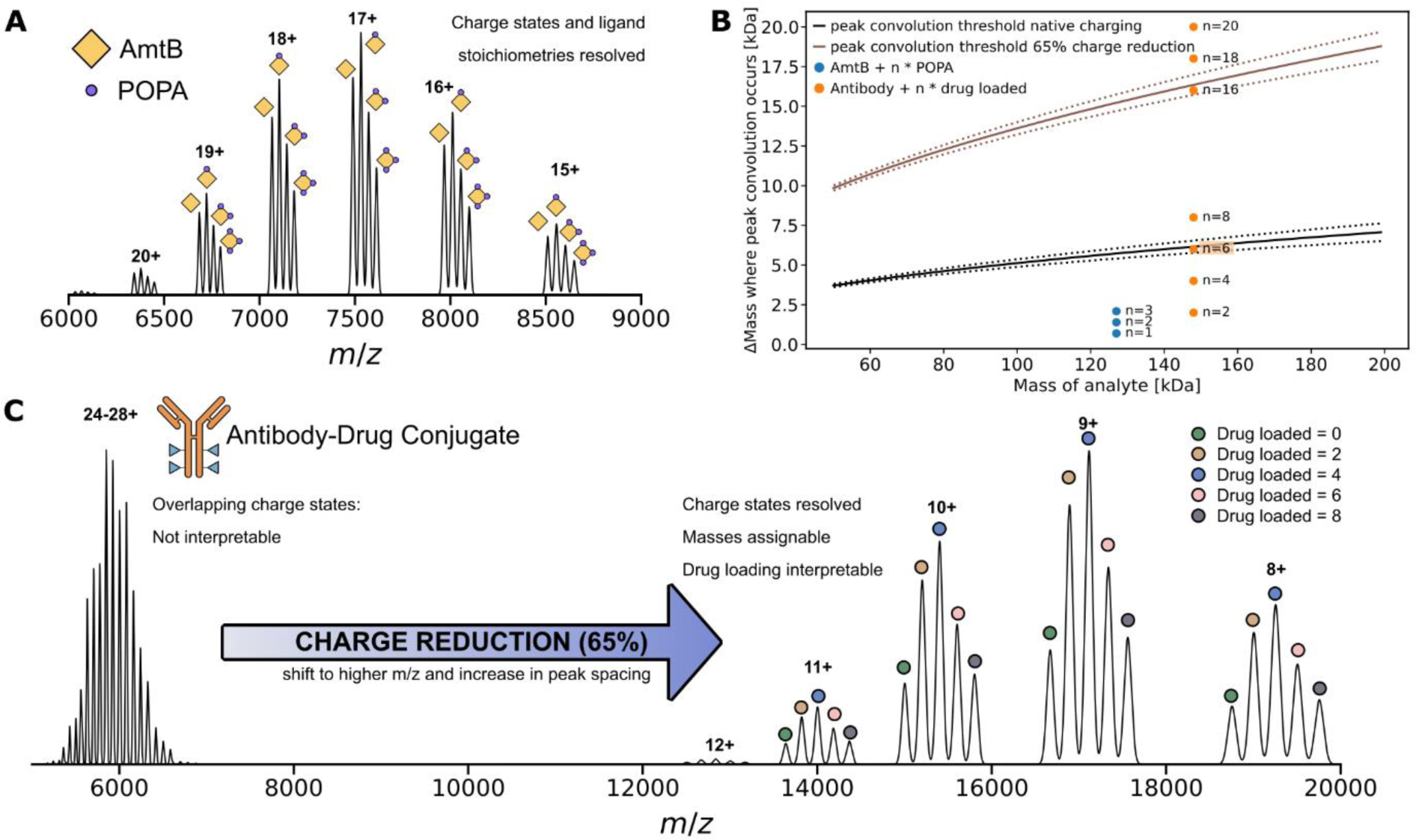
Simulations illustrating how charge reduction can deconvolve overlapping charge state distributions in native mass spectrometry. **A)** simulated native mass spectrum of ammonium transporter AmtB, a membrane protein for which charge states and lipid adducts (POPA) can be resolved without charge reduction, as an example of the ideal case in native mass spectrometry. Spectrum is modelled on the basis of spectra reported in the literature (*21*). **B)** Predicted overlap, or convolution, of peaks for ligand binding of AmtB and covalent modifications of an antibody (blue and orange dots). Depending on the Δmass of modifications, the z+1 charge state of (analyte mass + Δmass) and the z+ charge state of analyte mass will overlap when the corresponding Δmass value lies in the zone between the dotted lines, as is the case here for the n = 6 drug-loaded antibody. Black lines were calculated on the basis of equation 2 and assuming native charging behaviour (see also Suppl. Fig. 1), and brown lines are based on moderate charge reduction (assuming 65% charge reduction on average). Solid lines indicate the Δmass where exact signal overlap will arise while dotted lines bound a window of unresolvable peak overlap due to instrument limitations (here modelled for an Orbitrap analyser at resolution setting 1500, as a low resolution example). Coloured dots indicate positions on the chart of example native proteins (AmtB, blue; antibody drug-conjugate, orange) with ligand mass modifications (n = number of ligands). For the ADC there will be unresolvable convolution between adjacent charge states of the unmodified protein ion and the n = 6 drug loaded ions while n = 8 can still be resolved; moderate charge reduction resolves this by moving the peak convolution threshold to higher Δmass. **C)** simulated native mass spectrum of an antibody-drug conjugate showing overlapping charge states (signal around 6000 m/z) with native charging, precluding interpretation; charge reduction increases the separation between adjacent charge states, enabling charge state resolution and inference of the DAR (*20*). Simulations are modelled on the basis of real spectra reported in the literature (*20–22*).

Assuming the average native charge at a given mass (Suppl. Fig. 1), we can simulate how *Δmass* scales with the analyte mass, plotted as the solid line black line in Fig. 1B. The resolving power of the mass analyser furthermore adds a window of overlap to *Δmass* (see dotted lines in Fig. 1B), as for two peaks to be resolved by the instrument the following equation needs to be fulfilled: *m*/*z*_*ion*;2_ − *m*/*z*_*ion*;1_ > 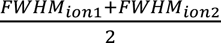. For the ammonia transporter AmtB and its POPA (1-palmitoyl-2-oleoyl-sn-glycero-3-phosphate) lipid ligands, it can clearly be seen in Fig. 1B that at its given native charging, all ligand-bound states lie well below this *Δmass* region where peak convolution occurs, no overlapping charge states are present, and ligand peaks are well resolved in Fig. 1A. However for an antibody drug-conjugate (ADC) with different numbers of drugs loaded, native charging and the larger size of loaded drugs predict that the ions of the two neighbouring drug-to-antibody ratio (DAR) variants 0 and 6 will fall within the peak overlap window and will be convolved (i.e., overlap) on lower resolution mass analysers, as reported experimentally (*20*). This behaviour will result in mixed charge state distributions which are often elusive for mass determination and quantification as it is not apparent from the convolved signals how many ion species are hiding within one peak (Fig. 1C and Suppl. Fig. 2).

The effect of charge reduction can also be simulated using equation 2, to estimate at which drug loading level (*Δmass*) peak convolution will appear again (Fig. 1B). In addition to ADCs, charge reduction has been exploited for separation of convolved signals and has assisted with characterization of for example, vaccine components, synthetic polymers and membrane proteins (*17, 20, 23–28*). For modelling electron capture charge-reduction, equation 2 should strictly be adapted to account for the mass of one additional Dalton per reduced charge, however this additional mass is negligible for the kilodalton and megadalton sized analytes discussed herein.

For the example in Fig. 1B, we have considered only a single charge state (the assumed average native charge state). Where other charge states are present in the spectrum (which have a different *Δmass* for peak convolution) it may be possible to make assignments from these other charge states; however heterogeneity from, for example, solvent adducts or glycoforms (in the case of ADCs) may make this more difficult. Furthermore, as analytes become heavier and/or multimeric, heterogeneity can also arise from the presence of many different types of ligands required for their biological functions, although these can often still be resolved with high instrument resolution or the use of chemical charge reduction (*29*). A further level of complexity can arise in the megadalton scale with analytes such as virus-like particles and nanocages, which can have high numbers of subunits and large encapsulated cargo molecules. Stochastic assembly of different subunits and/or variations in cargo loading can result in extreme heterogeneity in such systems, which can be beyond the capability of existing mass analysers to resolve. For example, adeno-associated virus (AAV) capsids are particles in the megadalton range used as vectors for gene therapy, and are an example of a highly heterogeneous sample (*22, 30*). The 60-mer AAV capsids (∼3.7 MDa) assemble stochastically from three structurally interchangeable subunit variants of different mass (VP1, VP2, VP3), resulting in an extremely heterogeneous mass distribution of 1891 possible capsid stoichiometries (Suppl. Fig. 3A) (*22*). Applying equation 2 with the mass and expected charging of such particles, it can be seen that the expected *Δmass* where peak convolution appears is in the range of about 21±9 kDa, in which peak overlap will occur at resolution settings reported for experimental spectra of these particles using Orbitrap™ mass analysers (*22*). This mass difference coincides with the mass difference of a single VP1 to VP3 substitution (21.9 kDa for AAV8) explaining the complex interference pattern which adeno-associated viruses display when charged natively (*22*). This complexity precludes the assignment of individual charge states to the convolved peaks, which is necessary for mass determination. Expansion of equation 2 to consider convolution of more distant charge states z_n_ and z_n+x_ yields the more general equation 3:

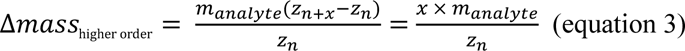

This shows for example that additional convolution of z_n_ with z_n+2_ or z_n+3_ will occur at *Δmass* two or three times that for z_n+1_. This leads to even greater convolution at integer multiple *Δmass* for extremely heterogeneous samples such as AAVs (Supp. Fig. 3C). Therefore in order to resolve the true charge state distribution of such particles, the peak convolution threshold has to be increased to much higher values than is possible with chemical charge reductions agents in solution, which only provide moderate charge reduction (see Suppl. Fig. 3B).

Alternative methods, involving manipulation of the analyte charge in the gas phase, have the potential to provide greater and more tuneable charge reduction; these can involve direct interaction of the ions with free electrons (electron capture) or gas-phase reactions in which electrons or protons are transferred between the protein and an electron donor or proton acceptor reagent (*31–33*). Charge reduction of natively folded proteins was in fact observed initially as an undesirable non-dissociative side-reaction of top-down fragmentation experiments using electron capture dissociation (ECD) or electron transfer dissociation (ETD) (*32, 34–37*). The full mechanistic picture of electron capture/transfer charge reduction has not yet been elucidated, but one idea is that peptide backbone fragmentation does actually occur as a result of the absorbed electron, but the extensive non-covalent interactions in the natively folded, now charged reduced precursor holds the fragments together tightly, such that it can only be dissociated upon further activation (*38–40*). Lermyte et al. showed that ETD could be used to achieve extensive charge reduction of native proteins to ∼120,000 *m/z* on a quadrupole-time of flight (Q-TOF) mass spectrometer, which is the effective mass range of this particular instrument (*39*). Charge reduction to even higher *m/z* scales offers potential for improved resolution of even greater heterogeneity in protein analytes, and can also serve as a test to explore and further develop the high *m/z* capabilities of existing instrumentation.

ECD was previously limited to Fourier Transform-Ion Cyclotron Resonance (FT-ICR) instruments (*41–44*). The recent development of electron capture cells, which use magnetic fields to trap an electron cloud in the path of the ion beam, made ECD compatible with commercial Orbitrap and Q-TOF instrument platforms and has greatly increased its practicality resulting in a recent flurry of research applying ECD for protein fragmentation and dissociation (*45–58*). The application of this new generation of ECD devices to charge reduction however remains neglected. In this work we explored the use of electron capture charge reduction (ECCR) using the ExD TQ-160 (e-MSion Inc.) electron emission cell on the Thermo Scientific™ Q Exactive™ UHMR (Ultra-High Mass Range) Orbitrap mass spectrometer (MS) (*45*). The UHMR MS is a popular instrument for native mass spectrometry due to its high resolving power, *m/z* range (specified up to 80,000 *m/z*) and enhanced activation and desolvation capabilities (*6, 59–61*). It has been successfully used for the characterization of megadalton complexes, such as ribosomes, hepatitis B virus capsids (3-4 MDa) and the flock house virus (∼9.3 MDa) (*59, 62*). Growing interest in the characterization of larger particles such as exosomes, the bacteriophage T5 (∼105 MDa), adenoviruses (up to 156 MDa) and carboxysomes (> 300 MDa), which would natively charge (Suppl. Fig. 1) above the 80,000 *m/z* specification of the UHMR MS, prompts the need to further explore the high *m/z* detection capabilities of existing Orbitrap instrumentation (*63–68*). We therefore set out to explore the use of the e-MSion ExD cell for ECCR on the UHMR with the following goals: (i) examine the tuneability of ECCR using the ExD cell, (ii) investigate the instrument parameters affecting the extent of charge reduction, (iii) use the high *m/z* ions thus generated to determine (and if possible extend) the upper *m/z* range of the UHMR, and (iv) explore the application of ECCR for the resolution of overlapping signals in extremely large, heterogeneous analytes.

## Results

### ECCR is tuneable and achieves up to 90% charge reduction of GroEL

In the modified QExactive UHMR MS, the ExD cell is positioned in the ion beam path after the quadrupole and replaces the transfer multipole (Suppl. Fig. 4). It consists of an electron-emitting coiled filament and seven additional lenses and lens magnets along the ion beam path (Fig. 2B) (*45*). When a current is passed through the filament with a voltage offset applied between it and lens 4, low energy electrons are emitted and radially confined by the magnetic fields of lens magnets 3 and 5. Application of negative voltages to the outer lenses confines the electrons axially. Electron density is limited primarily by space charge effects from other electrons, and the electrons move rapidly from the filament into the cell. A cloud of low energy electrons is consequently confined along the ion beam path. Tuning of the filament current and ExD cell voltages provides control over the electron capture process by controlling the distribution and density of the electron cloud within the cell, as well as affecting ion trajectories before, during and after interaction with the electron cloud.

**Figure 2.**
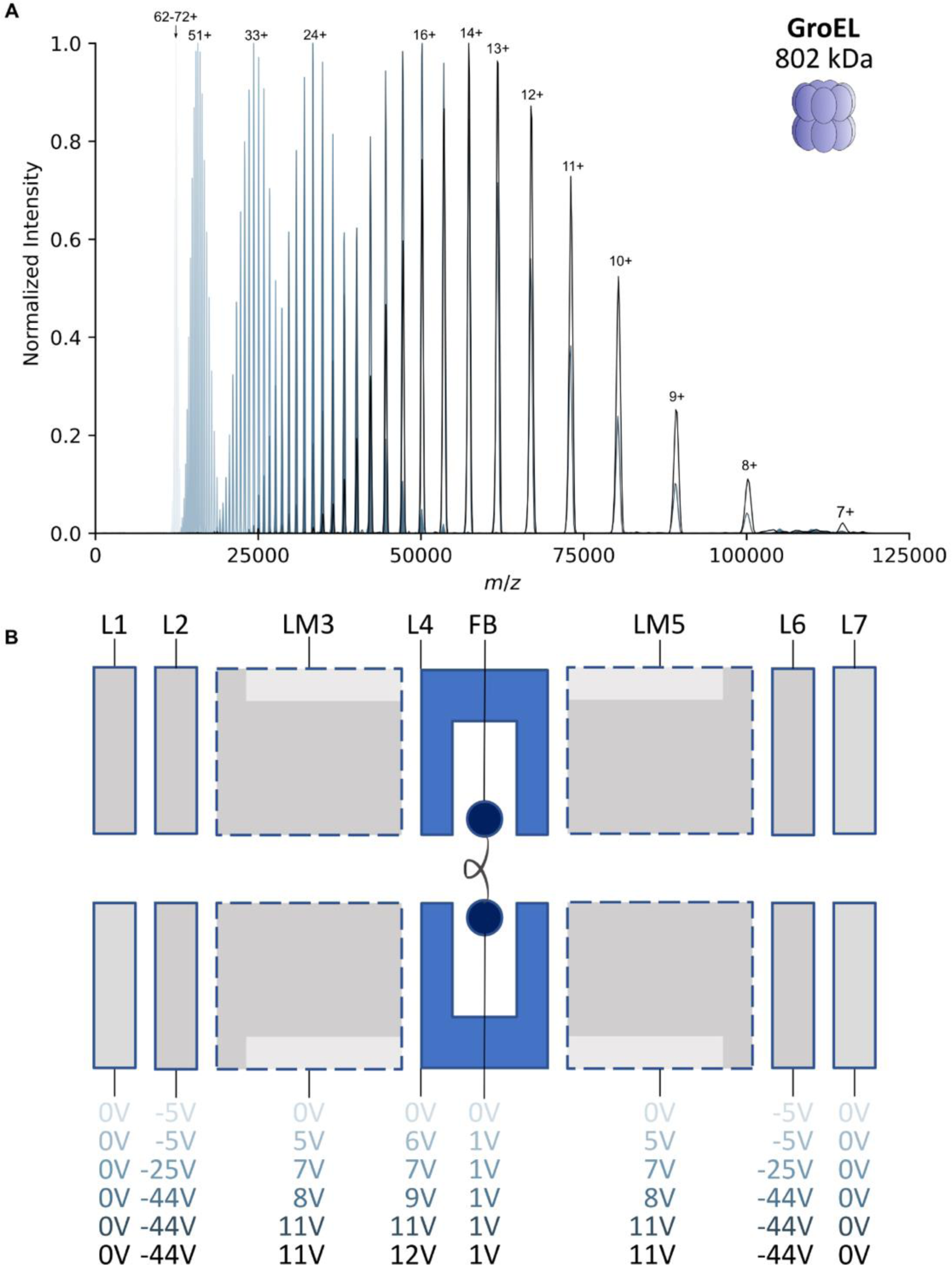
Stripping GroEL of 90% of its charge by ECCR. **A)** Native mass spectra of GroEL acquired under a normal native charging regime or with different amounts of electron capture charge reduction. Spectra were acquired with identical UHMR settings and spray conditions, but with different voltage settings on the ExD cell. **B)** Schematic of the ExD cell comprising lenses (L1-7), lens magnets (LM3 and LM5) and a metal filament (FB; filament bias voltage). Voltages applied to the elements of the ExD cell in order to obtain the spectra displayed in **A** are shown below the schematic, with colours matching those shown in part **A**. For the native charging spectrum (pale blue) the filament current was set to 0 A (i.e. ECD off), for all other spectra this was set to 2.3 A.

Using the 802 kDa bacterial chaperonin GroEL as a test case, tuneable charge reduction over a broad m/z range was achieved by varying the ExD voltages (Fig. 2) with fixed UHMR instrument settings. With the ExD cell tuned to a transmission only mode, without electron emission, GroEL was detected centered on its 65+ charge state at 12,340 *m/z*, with an observed native charge state envelop between 59+ and 70+ (*69*). −5 V was applied to L2 and L6 during this acquisition to assist with ion transmission. ECCR was initiated by applying the filament current (2.3 A), allowing it to stabilize, then setting the filament bias (FB) to 1 V and increasing the L4 voltage to 6 V in 1 V steps; resulting in electron emission. LM3 and LM5 were slowly increased to 5 V. These settings resulted in a clear and rapid shift of the entire GroEL charge state envelope to the right (Fig. 2A), light grey spectrum).

The amount of charge reduction could be controlled, and gradually increased by slowly increasing the voltages on L4, LM3 and LM5. It was observed consistently that greater ECCR results in a decrease in observed total ion signal, which has multiple causes. The intensity of the image current generated on the Orbitrap detector electrodes by an ion packet is linearly proportional to the total charge of the ion packet. Charge reduction will therefore cause signal intensity loss as a matter of instrumentation principles. Additional losses may arise from perturbation of ion trajectories by the electron cloud, lens voltages and from the electron-capture dissociation processes. It is therefore important to try to mitigate the loss of signal by adjusting settings to maximise transmission while also attaining the desired charge reduction. While increasing positive voltages on LM3, L4 and LM5 it was found that larger compensatory negative voltages applied to L2 and L6 were necessary to try to mitigate reduction in the total ion signal.

With maximal charge reduction (before reaching the limit of unacceptable signal loss) it was possible to charge reduce GroEL down to the 7+ charge state at 114,600 *m/z* (Fig. 2A, black spectrum), beyond the commercial specification of 80,000 *m/z*, without further instrument modifications. Charge states lower than 6+ were not observed, but charge reduction could be arbitrarily and easily achieved across the entire commercial *m/z* range and beyond, down to the 7+ charge state through tuning of the ExD cell voltages.

### Ion kinetic energy affects the extent of electron capture charge reduction

Variation in UHMR ion optics settings also influence the extent of charge reduction, especially settings which influence ion kinetic energy before the ions reach the ExD cell. This allows further control over the extent of charge reduction, by varying the transit time of the analyte ions through the electron cloud. To demonstrate this effect we varied the voltage offset experienced by the ions as they pass from the injection flatapole to the bent flatapole, prior to entry into the quadrupole and ExD cell (Fig. 3A). Ions accelerated by a greater voltage drop will acquire greater kinetic energy. Considering the 65+ charge state of GroEL in the high kinetic energy regime (Fig. 3A) with a DC voltage drop from 12 V to 2 V we can estimate the upper initial kinetic energy acquired by the ions as 650 eV neglecting collisions, and for the lowest kinetic energy regime (4 V to 2 V) as 130 eV, resulting in an up to five fold difference between the highest and lowest acquired kinetic energies in the experiment, and consequently up to √5 = 2.24 times different velocities. Maintaining fixed settings on the ExD cell and the rest of the instrument, we consistently found that spectra in the lower kinetic energy regime, when ions are slower, exhibited greater charge reduction (Figs. 3B and 3C). In native top-down ETD experiments using gas phase electron transfer reagents, it has been shown that increasing the reaction time leads to more extensive charge reduction (*40*). Our observation that reducing ion kinetic energy leads to more extensive electron capture charge reduction is consistent with these findings for ETD, as the slower-moving, lower-kinetic energy ions inherently spend more time in the ExD cell interacting with the electron cloud, increasing the opportunity to undergo electron capture events. It is interesting to note that, in contrast to the results of Fig. 2A, where tuning ExD voltages to increase charge reduction shifted the entire charge state distribution to lower charge states, tuning of the flatapole voltages instead extends the distribution to lower charge states, while maintaining a population of higher charge states. The origin of these distinct behaviours is not clear, but might be related to a larger diversity of pathways and kinetic energies of ions when flatapole acceleration is used. Nevertheless, the finding that control of ion kinetic energy also allows tuning of charge reduction is fruitful, as it provides an additional means to control and achieve more extensive charge reduction in combination with the voltage settings on the ExD cell.

**Figure 3.**
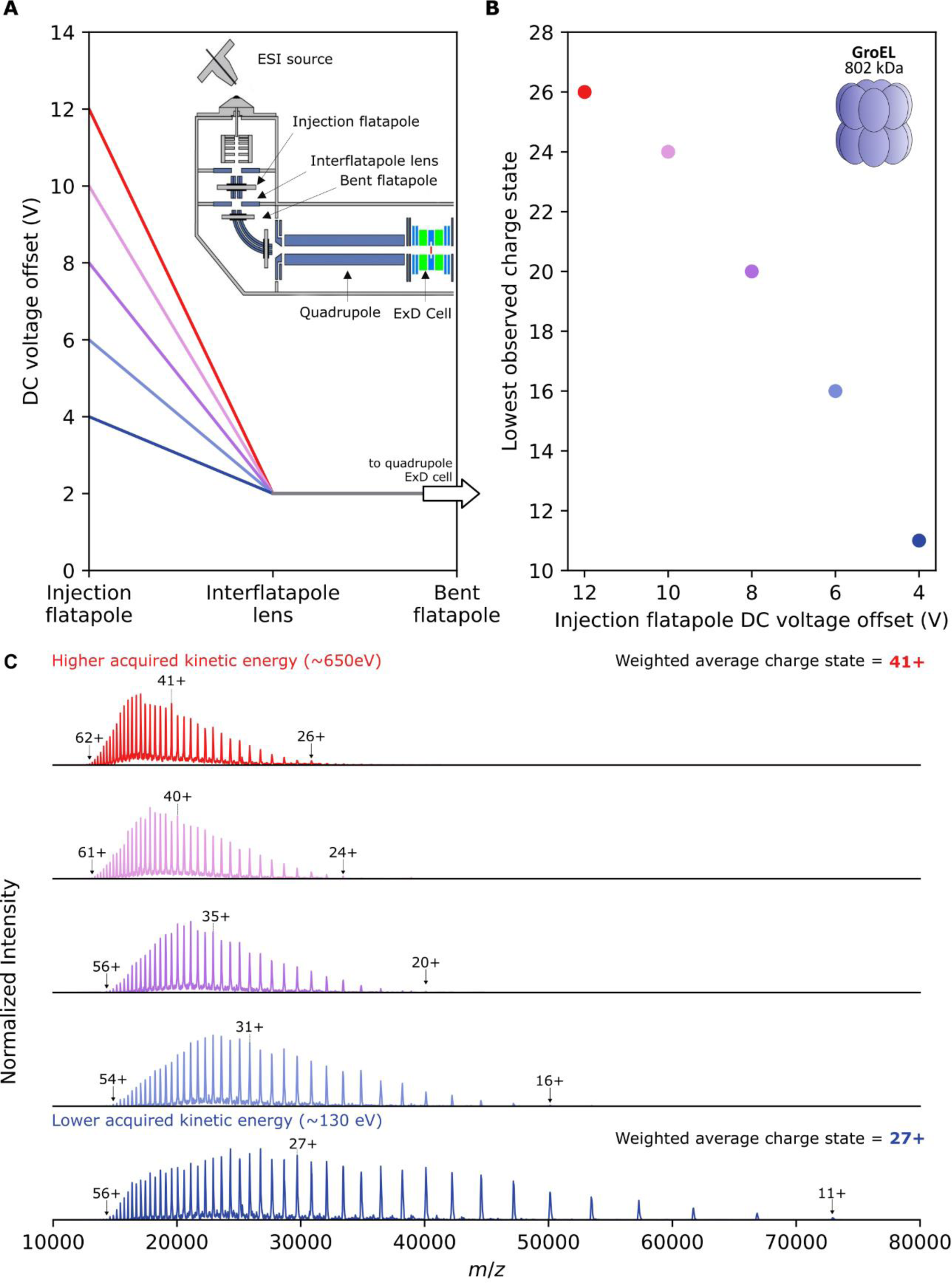
Ion kinetic energy affects the extent of charge reduction. **A)** Illustration of how the DC offset voltages of the injection flatapole can be varied (red: 12 V, through to dark blue: 4 V) while maintaining a fixed DC offset voltage of 2 V on the interflatapole lens and bent flatapole in order to control the kinetic energy of the ions as they enter the quadrupole and pass through to the ExD cell. The ions will acquire greater kinetic energy when the voltage offset is higher, and move faster through the ExD cell. Inset: schematic of the source and high vacuum region of the UHMR, with relevant ion optics labelled. **B)** Lowest detected charge state of GroEL observed during ECCR MS with fixed ExD cell voltages while varying the flatapole voltages as shown in panel **A**. **C)** The corresponding spectra for the data points in panel **B**, obtained with the flatapole settings shown in panel **A**.

### Simulating the effect of charge reduction on heterogeneous adeno-associated virus assemblies

Having demonstrated the feasibility of extensive ECCR on the UHMR instrument and identified the factors allowing us to control the extent of charge reduction, we next endeavoured to investigate its applicability to considerably larger and more heterogeneous analytes. An incidental advantage, when performing ECCR on heavier analytes, is that their larger exposed surface means each particle will acquire a greater number of charges during native nano-ESI (*11*), meaning they will induce greater signal per ion on the Orbitrap detector when charge reduced to the same *m/z* range as a smaller native protein with fewer charges. To this end we selected adeno-associated viruses (AAV) as suitably heavy (∼3.7-4 MDa) and extremely heterogeneous analytes.

AAV capsids are comprised of 60 subunits drawn from three structurally interchangeable proteins of different mass: VP1, VP2 and VP3. The prevailing model of AAV capsid assembly is that they assemble stochastically from the available expression pool of VP1, VP2 and VP3; producing a mixture of capsids of different VP stoichiometry and therefore mass (*22*). As a consequence of this mass heterogeneity (Suppl. Fig. 3A), under normal charging conditions, AAV native mass spectra produce an interference pattern from many overlapping signals of different mass and charge, as simulated in Fig. 4A. In the natively charging simulation there are for example up to 12 unique charge states (and multiple masses per charge state) overlapping within a single convolved peak. Such a spectrum eludes charge state assignment and therefore mass deconvolution. As noted in a recent publication, moderate charge reduction (simulated in Fig. 4A), for example by addition of TEAA, in fact appears to worsen this problem - producing even greater interference of misaligned charge states (*30*). Our simulations (Fig. 4A) predict however, that when AAVs are extensively charge reduced, the same charge states (of different masses) eventually all begin to group together with sufficient separation from adjacent charge states, such that charge state assignment should become possible beginning from around the 36+ charge state at ∼100,000 *m/z*. Correct charge state assignment should then enable calculation of the mean mass of the AAV capsid assemblies, a feat which has previously only been possible by alternative techniques such as charge detection mass spectrometry or mass photometry (*22, 70–73*).

**Figure 4.**
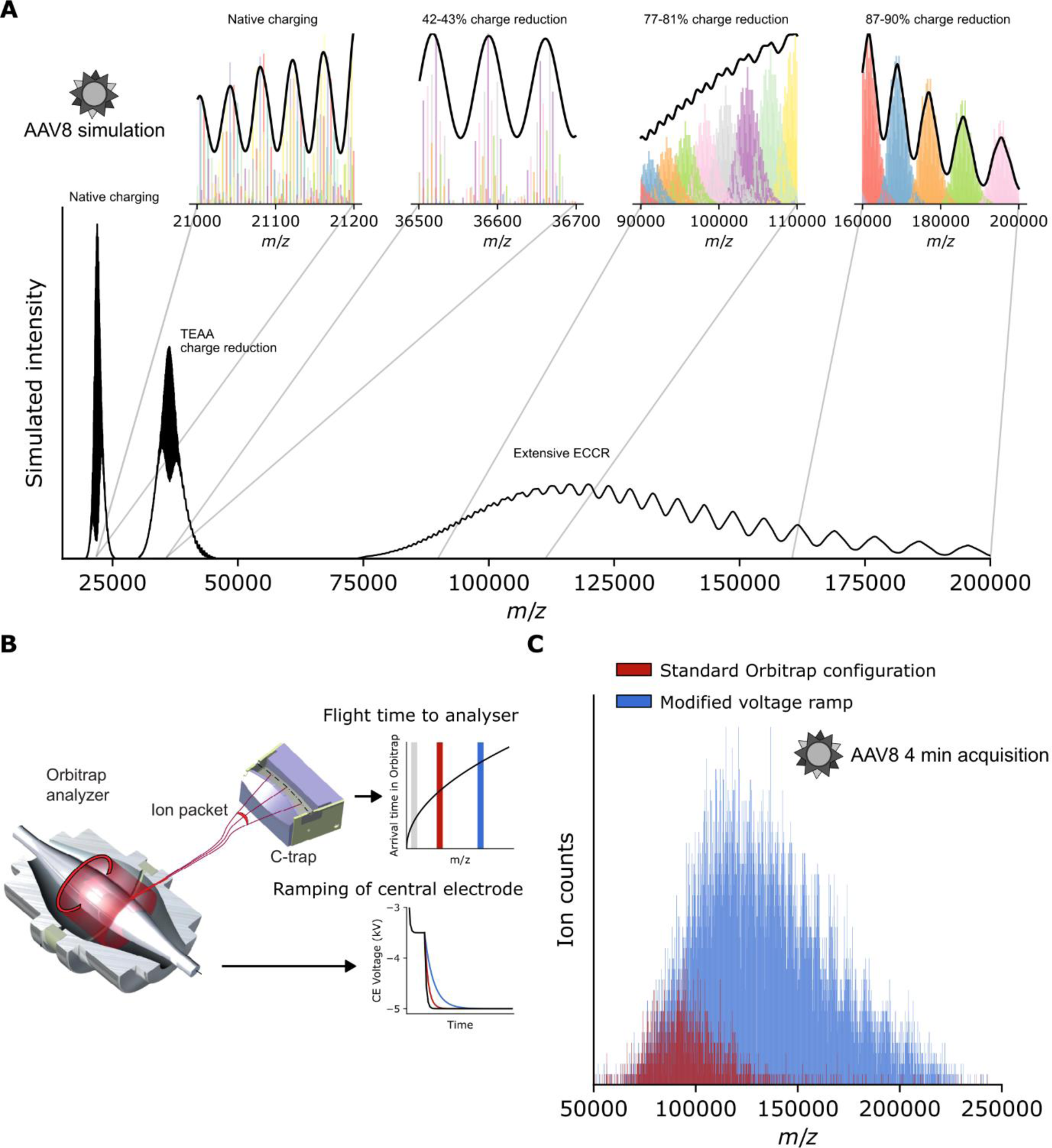
Charge state resolution of AAV8 virus samples should be achievable if charge reduced beyond 100,000 m/z; modification of the rate of the Orbitrap voltage ramp extends the accessible m/z range to 200,000 m/z and beyond. **A)** Simulated AAV8 mass spectra assuming stochastic capsid assembly from a pool of VP1, VP2 and VP3 subunits in 1:1:10 ratio. Simulations are shown for native charging conditions, charge reduction through the use of TEAA, and charge reduction above 100,000 m/z by ECCR at a resolution setting of 3000. Insets show simulated peak positions for all 1891 possible capsid stoichiometries, for charge states 166+ to 181+ (left) or 24+ to 39+ (right). Under native charging conditions, the numerous overlapping charge states produce a complex interference pattern. Aligned charge states become resolvable following charge reduction above 100,000 m/z. **B)** Illustration of the time-of-flight effect of ion injection from the C-trap into the Orbitrap analyser, and ramping of the voltage on the Orbitrap central electrode. **C)** Charge reduced AAV8 mass spectra (short 4 minute acquisitions) acquired with the standard UHMR configuration (red) or with the voltage ramp modification (blue) which enhances the detection capability above 100,000 m/z.

### Modifying the rate of the Orbitrap central electrode voltage ramp extends the upper *m/z* range to 200,000 *m/z* and beyond

Following the same methodological principles used to charge reduce GroEL, we were able to charge reduce empty AAV8 capsids to approximately 120,000 *m/z* (Fig. 4C, red spectrum), demonstrating that it is possible to reach the *m/z* region in which simulations indicate it should be possible to achieve charge state resolution. We noted with interest however that despite the greater charge (and therefore greater signal intensity) of the AAV8 capsids, the signal intensity faded out around 120,000-130,000 *m/z*, just like we observed with GroEL (Fig. 2A). This observation points towards instrumentation as the factor limiting the effective *m/z* range, especially considering that the maximally charge reduced spectra are acquired at *m/z* values 25% higher than the instrument specification. We therefore sought to increase the detection capabilities of the instrument above 100,000 *m/z*.

Injection of ions from the C-Trap into the Orbitrap analyser is a time-dependent process, with higher *m/z* ions arriving at the Orbitrap later than lower *m/z* ions (Fig. 4B). Meanwhile during injection, the Orbitrap central electrode voltage is ramped down from an initial injection voltage to the final negative measurement voltage (*74*). This ramping towards increasingly negative voltage compresses the ion trajectories, pulling them closer to the central electrode (electrodynamic squeezing), and is essential for the establishment of stable ion trajectories and avoidance of collision with the outer electrodes. Crucially, the timing of the voltage ramp needs to be aligned with the arrival time of the ions in the *m/z* range that is desired to be detected (*74*). Slowing the rate of the voltage ramp should increase the upper *m/z* limit by allowing more time for higher *m/z* ions to arrive at the Orbitrap entrance and adopt stable orbits by electrodynamic squeezing.

On a research-grade UHMR MS, we made modifications to the Orbitrap electronics to enable switching between the standard configuration and a configuration with a reduced rate of central electrode voltage ramping. With AAV8 charge-reduced as much as could be observed under the standard configuration (Fig. 4C, red spectrum) we observed that switching to the modified voltage ramp (Fig. 4C, blue spectrum) resulted in a dramatic increase in ion signals above 80,000 *m/z*, without a reduction in signal intensity below this threshold. The modified configuration furthermore enables detection of ions at *m/z* values which were not detected with the standard configuration, with clear ion signals being detected up to about 250,000 *m/z*. These are charge-reduced AAV8 capsid ions which were produced by ECCR and present in the instrument, but were simply of too high *m/z* to be detected in the standard configuration. Notably, resolved charge states are visible despite the short acquisition time.

### ECCR above 100,000 *m/z* enables charge state resolution and average mass determination of adeno-associated virus assemblies

Long acquisitions with the modified voltage ramp enabled definitive charge state resolution and average mass determination for different AAV8 preparations acquired from different sources (Fig. 5). For each preparation, measurements were conducted for both empty capsids and capsids filled with single-stranded DNA cargo encoding green fluorescent protein (GFP). The ECCR mass spectrum for preparation 1 of empty capsids exhibited a broad hump from ∼80,000 *m/z* to 200,000 *m/z*, bearing peaks to which charge states can unambiguously be assigned using the chevron method (see methods) (*75*). Calculation of the mass across all assigned charge states yields a mean mass measurement of 3.76 ± 0.03 MDa. This is consistent with a capsid stoichiometry of 1:1:10 (VP1:VP2:VP3; theoretical mass 3.74 MDa) and matches widely reported mass measurements for commercially available AAV8 preparations using alternative techniques, such as CDMS and mass photometry, confirming the correct charge state assignment (*8, 73*). For further confirmation of the mass we performed CDMS on the same sample (Fig. 5A inset). We observed consistent mass differences across the charge states within each of the ECCR spectra, arising from reduced desolvation of the more charge reduced ions; such ions experience reduced acceleration and therefore less collisional desolvation in the HCD cell. This mass difference assists in charge state assignment: the correct set of charge assignments being that which satisfies two constraints: (i) minimization of the standard deviation in mass across the charge state distribution (ii) an inverse correlation between charge and mass. This data represents the first ever deconvolution of the charge state distribution of extremely heterogeneous AAV assemblies by native mass spectrometry. AAV8 capsids loaded with genetic material could also be charge state resolved by ECCR (Fig. 5B). This is surprising, due the additional mass and heterogeneity added by the genetic vector, which can consist of either sense or anti-sense DNA. The mean mass difference between the full and empty particles points towards a genome mass of 0.900 ± 0.046 MDa, again consistent with literature and the diverse masses of such megadalton particles (*8*).

**Figure 5.**
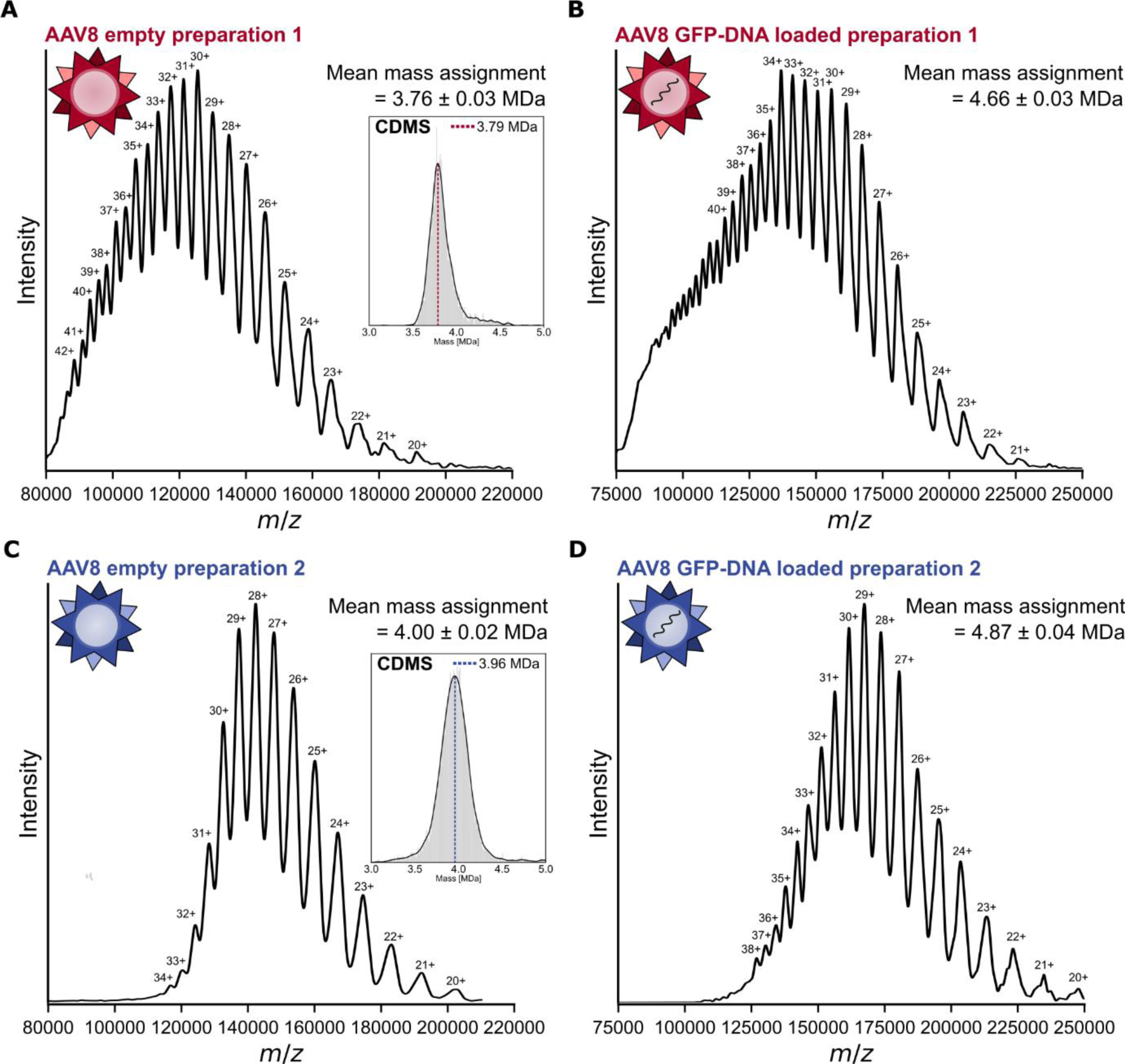
Charge state resolution and mean mass determination of different AAV8 preparations by electron capture charge reduction. Mass spectra of charge reduced AAV8 from two different preparations, enabling charge state assignment and mean mass determination. **A)** preparation 1 empty capsids, inset: charge detection mass spectrum obtained from the same sample. **B)** preparation 1 capsids filled with CMV-GFP genetic cargo. **C)** preparation 2 empty capsids, inset: charge detection mass spectrum obtained from the same sample. **D)** preparation 2 capsids filled with GFP encoding DNA vector.

AAV preparations from different suppliers and expression systems are known to exhibit variation in stoichiometry and genome packaging and therefore in mass (*73*). We therefore performed ECCR on a different preparation (preparation 2) of the same AAV8 serotype, again with empty capsids and capsids filled with genetic material (Fig. 5C and 5D). These samples were charge reduced slightly more than the samples for preparation 1 under similar conditions, and once again ECCR yielded charge state resolution and enabled assignment. These spectra yielded a higher than expected mass for empty AAV8 (4.00 ± 0.02 MDa measured vs. 3.74 MDa theoretical mass assuming 1:1:10 VP1:VP2:VP3 ratio), but the greater mass is supported by CDMS (Fig. 5C inset) and a genome mass of 0.87 ± 0.04 MDa determined by comparison of the full and empty spectra. The additional mass may be due to higher than expected VP1:VP2:VP3 ratio or unexpected encapsulations.

The m/z range and charge reduction displayed in these spectra are remarkable. With ions detected up to 250,000 m/z, these represent the highest m/z ions ever detected on an Orbitrap mass spectrometer; furthermore the ions were detected over an m/z range of ∼150,000 m/z, which is almost twice as large as the upper m/z commercial specification of the UHMR. It is furthermore remarkable that these gas phase virus particles are able to capture ∼140 electrons, experience up to ∼90% charge reduction and survive intact until detected, without dissociation.

## Discussion/Conclusions

This work demonstrates that controllable electron capture charge reduction can be achieved over a broad m/z range on the Q Exactive UHMR mass spectrometer, with the charge-state resolution and therefore mean mass determination of highly heterogeneous stochastically assembling adeno-associated virus capsids as a capstone application. Charge-state assignment of AAVs had until now been impossible by conventional native mass spectrometry, and this had stimulated the development of alternative techniques such as charge detection mass spectrometry (*8, 70, 76*). Compared with charge detection mass spectrometry, charge state resolution via ECCR has the advantage of more confident charge state determination and has less stringent requirements in terms of Orbitrap gas pressure or ion stability than CDMS. It has recently been highlighted that beyond AAVs, stochastic assembly of macromolecular complexes may be more common than previously realized; a phenomenon which has been overlooked and can lead to erroneous charge assignments (*30*). While limited charge reduction can potentially worsen the spectral appearance of such spectra, our work shows that extensive charge reduction presents an elegant and straightforward solution even for extreme cases such as AAVs. Given the increasing use of viral and lipid nanoparticle-based delivery systems for vaccination and gene therapy, ECCR presents new possibilities for characterizing these often highly heterogeneous systems (*77–79*).

Decreased desolvation of extensively charge reduced species due to their lesser acceleration by electric fields is a major factor limiting resolution in these experiments however. Combining ECCR with charge state independent methods of desolvation, such as IR photoactivation, could greatly enhance the power of this technique and is a promising direction for future research (*80, 81*). In the fields of gas-phase chemistry and electron capture/transfer dissociation it is a remarkable feat that viral capsids in the gas phase can capture over 100 electrons without dissociation.

This work furthermore represents a technical advancement in Orbitrap instrumentation. Without modification, we have shown that the UHMR can transmit and detect ions up to ∼130,000 *m/z*; 50,000 m/z higher than its 80,000 *m/z* commercial specification. With modifications, we were able to detect ion signals at ∼250,000 *m/z:* the highest *m/z* ions recorded to date on an Orbitrap mass spectrometer. This positions Orbitrap native mass spectrometry well in the context of growing interest in the characterization of larger particles such as exosomes, adenoviruses and carboxysomes (*63–66*). The 156 MDa human adenovirus 5, for example, typically charges with charge states around 1000+ placing it at ∼156,000 *m/z* – well within the detection capability of our modified UHMR (*65*). At this *m/z* range, adenovirus would have approximately 50 times the number of charges than the 24+ charge state of adeno-associated viruses which appear in the same *m/z* region; the 156 MDa adenovirus would therefore produce 50 times more signal and be even easier to detect than the charge reduced AAV ions detected in this work. With the *m/z* range demonstrated by our modified UHMR, detection of ions up to 550 MDa in mass is theoretically possible assuming native charging behaviour (Suppl. Fig. 1).

The high *m/z* region and extensive charge reduction to this region presents a number of technically interesting features and applications. High *m/z* ions, having lower frequencies, travel reduced distances in the Orbitrap analyser at lower velocities compared to lower *m/z* ions over the same transient time (Suppl. Fig. 5), resulting in fewer and less energetic collisions and hence more stable trajectories (*62*). This has potential applications for use cases such as Orbitrap CDMS and ultra-long transient acquisitions, for which ion stability is a key factor in the quality of results. We have furthermore shown how extensive and tuneable charge reduction to an *m/z* range of choice can serve as a useful tool in instrumentation development, allowing assessment of instrument capabilities over its entire *m/z* range. The ability to generate ions at controllable *m/z* furthermore has potential application for *m/z* calibration and for charge detection calibration in CDMS.

Various terminologies have been used to describe the charge reducing effect of electron and proton capture/transfer, such as ECnoD (electron capture no dissociation), charge reduction ETD, and proton capture charge reduction. We propose that the research community adopts the terminology: **e**lectron **c**apture **c**harge **r**eduction (ECCR), **e**lectron **t**ransfer **c**harge **r**eduction (ETCR) and **p**roton **t**ransfer **c**harge **r**eduction (PTCR) when these techniques are used for charge reduction applications where fragmentation is not the objective. These new terminology and acronyms make clear which type of charge reduction process is being employed, and distinguish this application from fragmentation experiments involving ECD and ETD.

Given the exciting results demonstrated here, and the present practical availability of devices for generating and confining free electrons within commercial mass spectrometers, we believe that electron capture charge reduction (ECCR) is set to enable exciting progress in the resolution of convolved signals in native mass spectrometry, especially when combined with additional emerging techniques such as charge detection mass spectrometry, ultra-long transients and surface induced dissociation.

## Materials and Methods

### Preparation of protein samples for native MS

Recombinant *E. coli* chaperonin GroEL was purchased from Sigma Aldrich. For purification the lyophilized powder was dissolved at a concentration of 1 mg/mL in 1 mL of reconstitution buffer (20 mM tris acetate, 50 mM potassium chloride, 0.5 mM EDTA, 1 mM ATP and 5 mM MgCl_2_) and vortexed for 1 hour at room temperature. 200 µL of ice cooled methanol was added and the solution was vortexed again for 1 hour at room temperature. Assembled GroEL oligomer was precipitated by addition of 600 µL ice cooled acetone and left for four hours at 5 °C. The supernatant was removed and the precipitate redissolved in the reconstitution buffer. For native MS application, GroEL was buffer exchanged into ammonium acetate (50 mM) by overnight dialysis at 5 °C and used for native MS at ∼5 µM final concentration.

Preparation 1 empty and full (filled with CMV-GFP genetic cargo) AAV8 capsids were purchased from Virovek. Preparation 2 empty and full (filled with zsGreen GFP genetic cargo) AAV8 capsids were expressed and purified as described below. All AAV8 samples were buffer exchanged into ammonium acetate (50 mM) by overnight dialysis at 5 °C and used for native MS at ∼100 nM final concentration.

### AAV8 expression and purification

HEK Viral Production cells (ThermoFisher) were maintained in suspension in Freestyle F17 (ThermoFisher) chemically defined, serum-free media supplemented with 1× Glutamax (Gibco). The cells were cultured in 2 L roller bottles (Greiner) at 37 °C, 7% CO_2_ and 135 rpm agitation in a humidified atmosphere. The plasmids used were pAAV_zsGreen, pAAV_RC8, pAAV_Helper (produced in-house). Cells were seeded into 9 L medium at 0.5 × 106 cells per mL in a wave bioreactor (Cytiva) on a Wave25 platform. 24 hours post-inoculation the culture was triple transfected using PEIMax (Polyplus) and three plasmids at a ratio of 2 : 1.5 : 1. To produce empty AAV particles, the culture were transfected identically except with the absence of the transgene plasmid (pAAV_zsGreen).

Each culture was harvested by batch centrifugation (3800*g* for 15 min at 4 °C) 72 hours post transfection to separate the cell fraction from the supernatant. The resulting cell pellet was freeze-thawed thrice and re-suspended to 10% (w/v) in a lysis buffer containing 5% (v/v) Polysorbate 80 and 20 U mL^−1^ benzonase (Merck). After 1 h incubation at 37 °C, the lysate was clarified by depth filtration followed by sequential filtration through 0.8 and 0.45 μm filters. The clarified lysate was pooled with filtered supernatant and concentrated by TFF on an Ultracel Pellicon 2, 300 kDa MWCO cassette (Merck) before loading onto a pre-equilibrated POROS CaptureSelect AAVX affinity column (Thermo Fisher) at 1 × 1013 vg per mL of resin. After re-equilibration, AAV was eluted with a low-pH buffer and immediately neutralised with 1 M tris base. Neutralised eluate was concentrated and buffer exchanged into PBS using an Amicon Ultra, 100 kDa MWCO concentrator (Merck) before storage at −80 °C.

### VLP expression and purification

The protein sequences used in the design of constructs for expression of AAV8 VP1, VP3 and Assembly-Activating Protein (AAP) proteins were derived from UniProt entry Q8JQF8. A 6×His tag was placed within an exposed loop of the VP1 protein, VP3 was left untagged and AAP was tagged at the N-terminus with the Flag tag. The VP1 coding region followed by an IRES and the AAP coding region was cloned into the pDEST12.2 OriP (Thermo Fisher) plasmid. The VP3 protein coding region was cloned into a separate pDEST12.2 OriP (Thermo Fisher) plasmid. The constructs were expressed in suspension Chinese Hamster Ovary (CHO) cells following transfection with a 1 : 2 molar ratio of VP1/AAP plasmid to VP3 plasmid and capsids were purified from the conditioned media using standard immobilized metal affinity chromatography, anion exchange and size exclusion chromatography purification.

### Native MS instrumentation

ECCR spectra were acquired on a modified, research grade, Q Exactive-UHMR instrument. The instrument was fitted with an ExD TQ-160 (e-MSion) electron emission cell, replacing the transfer multipole. The detector pulser board was physically modified so that the rate of the voltage ramp on the Orbitrap central electrode could be switched from the standard factory configuration to a modified configuration with reduced slew rate. 3 µL aliquots of sample were injected into gold and palladium coated borosilicate glass capillaries, prepared in-house with a model P-97 micropipette puller (Sutter Instrument Company) and sprayed into the instrument by nano-electrospray ionization.

### Native MS and ECCR of GroEL

The effect of ExD cell settings on ECCR was assessed with fixed DC voltage offsets of 5 V on the injection flatapole, 4 V on the inter-flatapole lens and 2 V on the bent flatapole, while varying voltages on the ExD cell as shown in Fig. 2B. For the native charging experiment, with the ExD cell in transmission only mode, the filament current was set to 0 A, for all other spectra this was set to 2.3 A. Data were acquired from the same sample-loaded nanoESI capillary, which sprayed continuously during the experiments.

The effect of ion kinetic energy on ECCR was assessed while maintaining a fixed set of ExD cell voltages (L1 = 0, L2 = −27, LM3 = 6.5, L4 = 6.5, FB = 1, LM5 = 6.5, L6 = −27, L7 = 0) and 2.3 A filament current. The interflatapole lens (2 V) and bent flatapole (2 V) were held at fixed DC offset voltages, while the injection flatapole lens was decreased from 12 V to 4 V in 2 V every 5 minutes as shown in Fig. 3. Data were acquired from the same sample-loaded nanoESI capillary, which sprayed continuously during the experiments.

Spectra were acquired using in-source trapping (IST, −100 V) without HCD cell activation. N_2_ was used as the trapping buffer gas with typical UHV pressure reading of 5.5 x 10^−10^ mbar. High m/z ion transfer target and high m/z detector optimization settings were used, with the standard (unmodified) configuration of the Orbitrap voltage ramp rate.

### Native MS and ECCR of AAV8 preparations

AAV8 samples in 50 mM ammonium acetate were spiked with 25 mM triethylammonium acetate immediately prior to nanoESI in order to achieve limited chemical charge reduction prior to ECCR. The preceding chemical charge reduction helped in achieving greater overall maximal charge reduction. To optimize charge reduction, desolvation and signal quality, ExD cell voltages and UHMR ion transfer and desolvation parameters were tuned for each sample and are presented in Suppl. Table S1.

Xenon was used as a collision gas in the HCD cell, with exemplary UHV vacuum pressure readout of 7.7 x 10^−10^ mbar. Ion transfer and detector optimizations settings were set to high m/z. Spectra were recorded with a 4096 ms transient time using magnitude mode Fourier transform with FFT enabled. Ions were detected as individual particles, similar to a CDMS experiment, due to low sample concentration, transmission losses from ECCR and the heterogeneity of AAV8 assemblies (*8*).

For the experiments shown in Fig. 3B, which assess the effect of the Orbitrap voltage ramp rate on detection of ultrahigh m/z species above 100,000 m/z, AAV8 preparation 1 empty was charge reduced as much as possible without unacceptable loss of ion transmission and a spectrum was acquired for four minutes using the standard Orbitrap configuration. The Orbitrap configuration was then rapidly switched to the modified configuration with reduced voltage ramp rate and electrospray was restarted from the same needle, in the same position, with all other instrument settings the same and acquired again for four minutes. The results were reproducible when switching back to the standard configuration. All other AAV8 spectra were recorded using the modified Orbitrap configuration and acquired for 1-5 hours depending on spray stability.

### Analysis of AAV8 ECCR mass spectra

Raw acquisition files were converted to mzXML file format using the MSConvert tool and vendor peak picking algorithm within the ProteoWizard Toolkit (*82*). The mzXML files were analysed using python scripts making use of the Pyteomics, NumPy, pandas, SciPy and Matplotlib packages (*83–88*). Electrical noise signals which were persistent across many scans were manually identified and removed, and a custom decaying noise filter function, which accounts for the decreasing intensity of lower charged ions, was used to filter baseline noise. A histogram with 5 m/z bin spacing was constructed from the filtered ion signals, smoothened by Savitzky-Golay filtering and peaks were picked using the peak picking function of SciPy (*87*). Charge states were assigned using the chevron method, to find the charge assignment which is closest to the minimum of the standard deviation of mean mass calculated across all peaks, and which correctly exhibits differences in desolvation at different charge states (lower charge state peaks should have mass > mean mass; higher charge state peaks should have mass < mean mass) (*75*). The correct charge state assignment was always found to be one charge higher than that which globally minimised the standard deviation.

### Charge Detection MS

CDMS spectra were acquired on a standard Q Exactive-UHMR instrument (ThermoFisher Scientific), without modification, following published protocol and analysis scripts (*8*). The charge-intensity constant was calibrated using GroEL as a reference calibrant. Transients were recorded in enhanced Fourier transform mode at 50k and 100k resolution setting and with 800 ms injection time.

### Simulation of AAV8 charge reduction

Published python scripts for the simulation of stochastic assembly of AAV capsids and the resulting heterogeneous native mass spectrum were adapted so that the spectrum could be simulated at different levels of charge reduction, and modelling the decreased desolvation of charge reduced species (*22*). The simulations furthermore model the expected m/z dependency of the resolution in an Orbitrap instrument. Simulations were performed for a VP1:VP2:VP3 expression ratio of 1:1:10 with masses 81667.3 Da (VP1), 66518.6 Da (VP2) and 59763.1 Da (VP3).

## Acknowledgements

KIPLH is funded by Biotechnology and Biological Sciences Research Council grant BB/M011151/1. We acknowledge the use of the Biomolecular Mass Spectrometry Facility (University of Leeds) and the assistance of Dr James Ault. The authors thank Dr Ralf Hartmann, Dr Dmitry Strelnikov, Dr Konstantin Aizikov, Dr Eduard Denisov, Dr Denis Chernyshev, and Dr Frederik Busse (Thermo Fisher Scientific) and Prof. Joe Beckman (Oregon State University) for helpful discussions during this work.

## Competing interests

The authors declare the following competing financial interests: Tobias Woerner, Maria Reinhardt-Szyba, Kyle Fort and Alexander Makarov are employees of Thermo Fisher Scientific, manufacturer of instrumentation used in this work.

## Supplementary Materials

**Suppl. Table S1.** UHMR and ExD cell tuning parameters used for acquisition of AAV8 ECCR mass spectra.

| UHMR and ExD cell tuning parameters | AAV8 preparation 1 empty | AAV8 preparation 1 filled | AAV8 preparation 2 empty | AAV8 preparation 2 filled |
| --- | --- | --- | --- | --- |
| Filament current | 2.3 A | 2.3 A | 2.3 A | 2.3 A |
| L1 voltage | 0 V | 0 V | 0 V | 0 V |
| L2 voltage | -36.2 V | -52.7 V | -26 V | -58.3 V |
| LM3 voltage | 5.7 V | 5.7 V | 5.6 V | 5.7 V |
| L4 voltage | 6.1 V | 6.7 V | 6 V | 7.1 V |
| FB voltage | 1.0 V | 1.0 V | 1.0 V | 1.0 V |
| LM5 voltage | 5.7 V | 5.7 V | 5.6 V | 5.7 V |
| L6 voltage | -36.2 V | -52.7 V | -26 V | -58.3 V |
| L7 voltage | 0 V | 0 V | 0 V | 0 V |
| In-source trapping voltage | -135 V | -150 V | -120 V | -150 V |
| HCD energy | 212 V | 212 V | 125 V | 185 V |
| Injection flatapole DC offset | 3.5 V | 3.5 V | 4 V | 4 V |
| Interflatapole lens DC offset | 2.5 V | 3 V | 3 V | 3 V |
| Bent flatapole lens DC offset | 2.0 V | 2.0 V | 2.5 V | 2.5 V |

**Suppl. Fig. 1.**
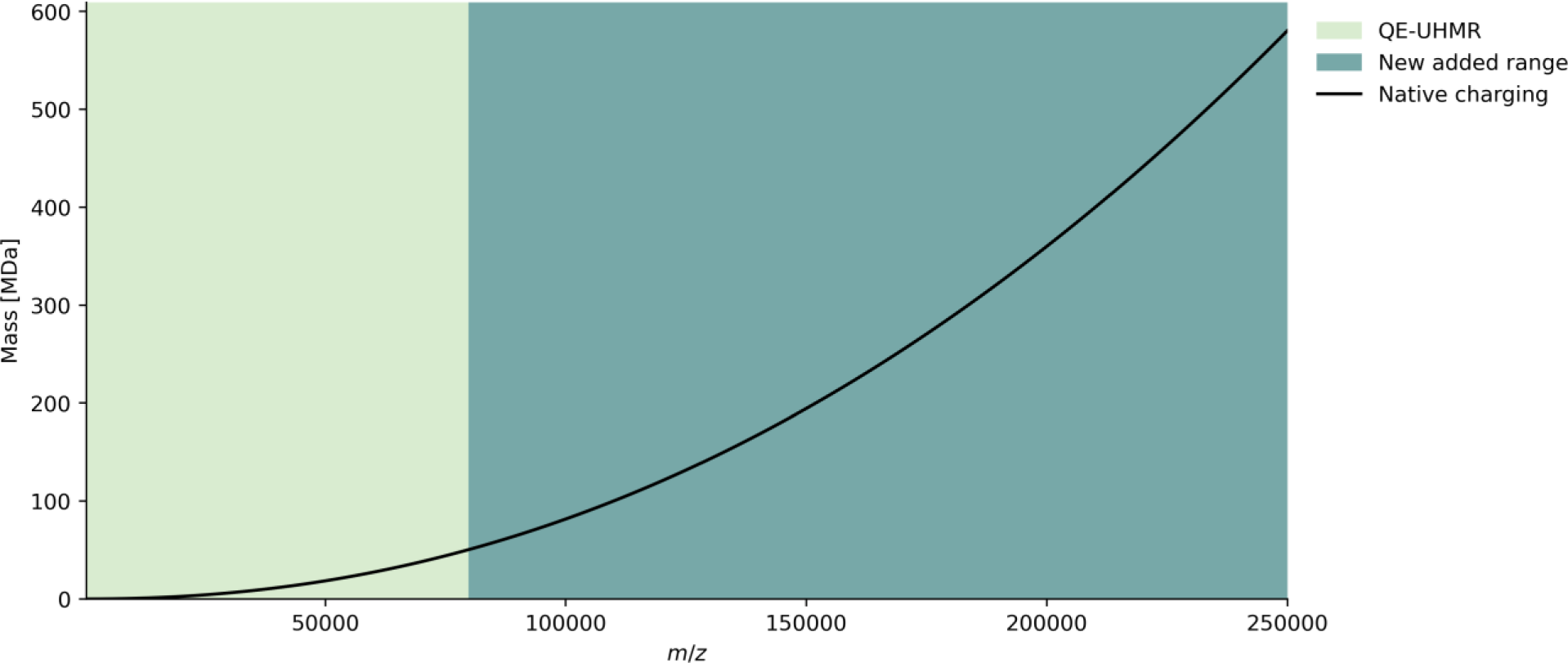
Expected average m/z position of biomolecular analytes assuming native charging and near-spherical shape and on the basis of empirical data (*7, 89*). The plot background is coloured to illustrate the standard m/z range of the QExactive UHMR MS and the further range newly added in this work.

**Suppl. Fig. 2.**
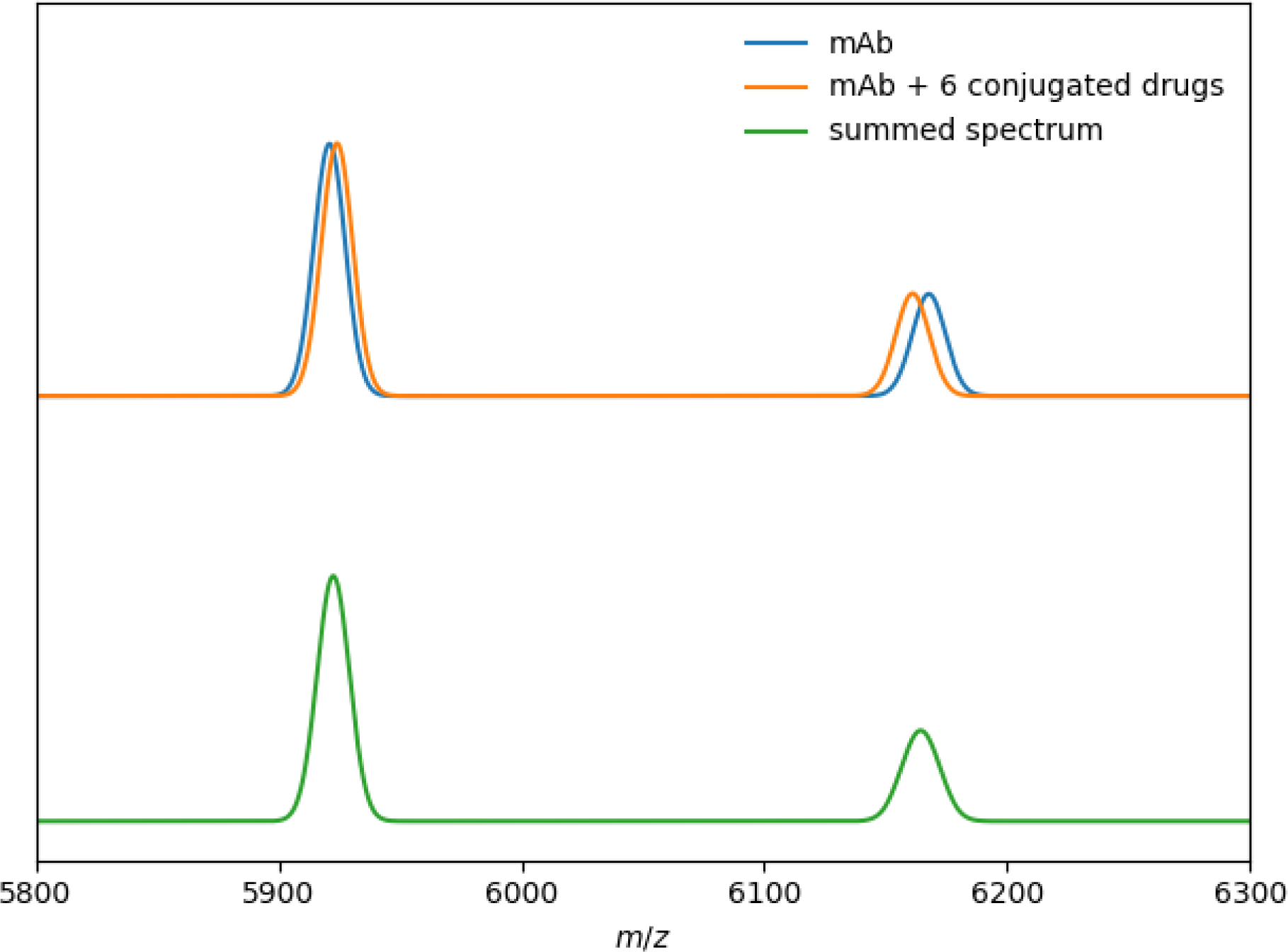
**(Supplement to Fig. 1).** Simulated peak overlap between the 26+/27+, and 27+/28+ charge states of a monoclonal antibody (mass = 148 kDa) and its 6 drug conjugated form (mass = 154 kDa) at an Orbitrap resolution setting of 1500.

**Suppl. Fig. 3.**
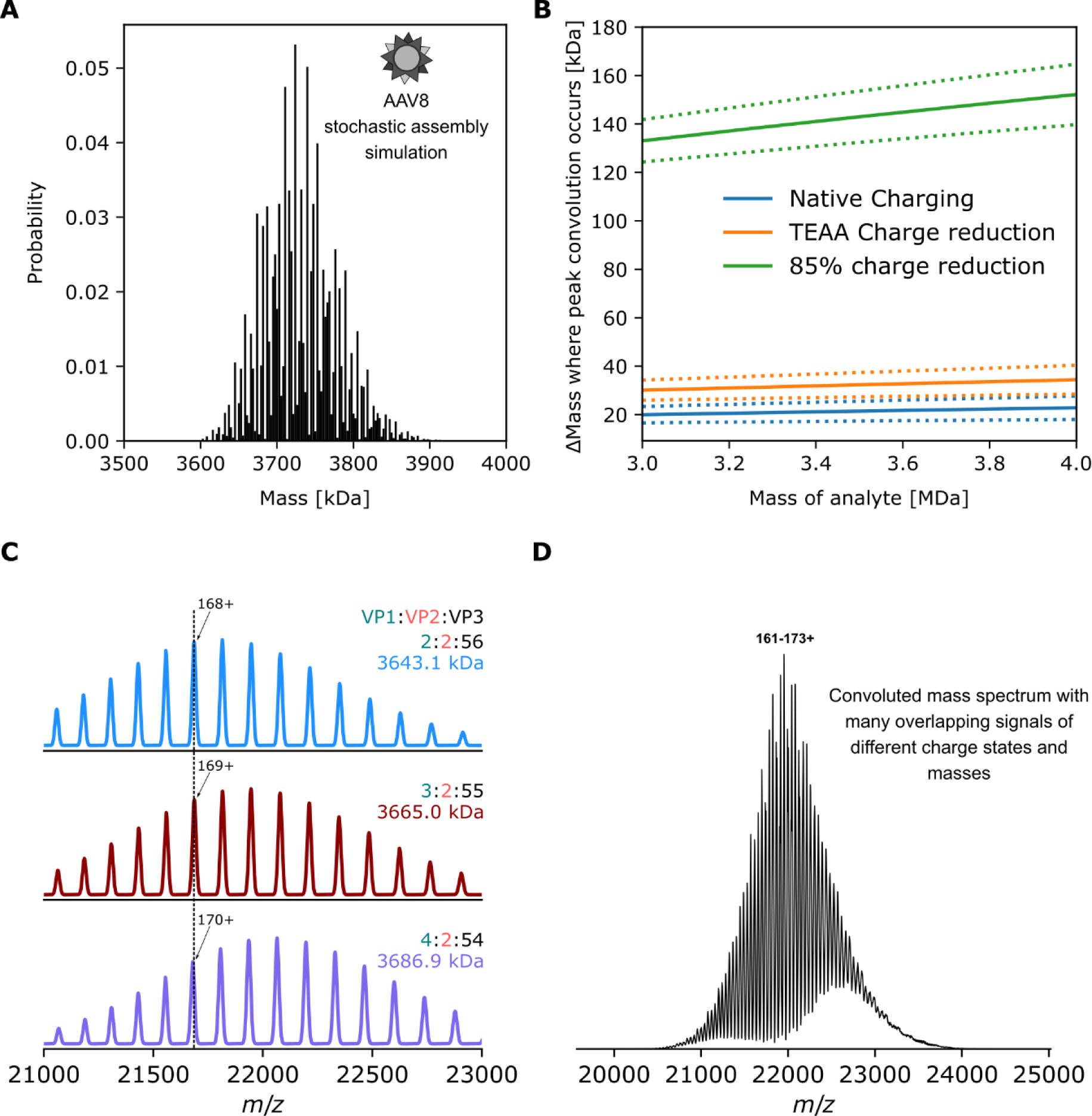
Simulated mass distribution, peak convolution and native mass spectrum of AAV8 capsid assemblies. **A)** probability distribution for 1891 possible capsid masses, assuming each capsid is assembled by 60 random draws from a subunit pool of VP1 (81667.3 Da), VP2 (66518.6) and VP3 (59763.1) in a VP1:VP2:VP3 population ratio of 1:1:10. **B)** Δmass of modifications for which there will be convolution between the z+1 charge state of (analyte mass + Δmass) and the z+ charge state of analyte mass, on the basis of equation 2 and assuming native average charging behaviour (blue), moderate charge reduction with TEAA (orange) or 85% charge reduction (green). Solid lines indicate the Δmass where exact signal overlap will arise, dotted lines bound a window of unresolvable peak overlap due to instrument limitations (here modelled for an Orbitrap transient length of 32 ms, as reported in the literature (*22*). **C)** Simulated native mass spectra for three pure capsid masses, illustrating coinciding signals of different charge and mass. **D)** Simulated native mass spectrum of the mixed capsid assemblies, with mass populations as shown in panel A. The overlap of many signals of different charge and mass produces a complex interference pattern.

**Suppl. Fig. 4.**
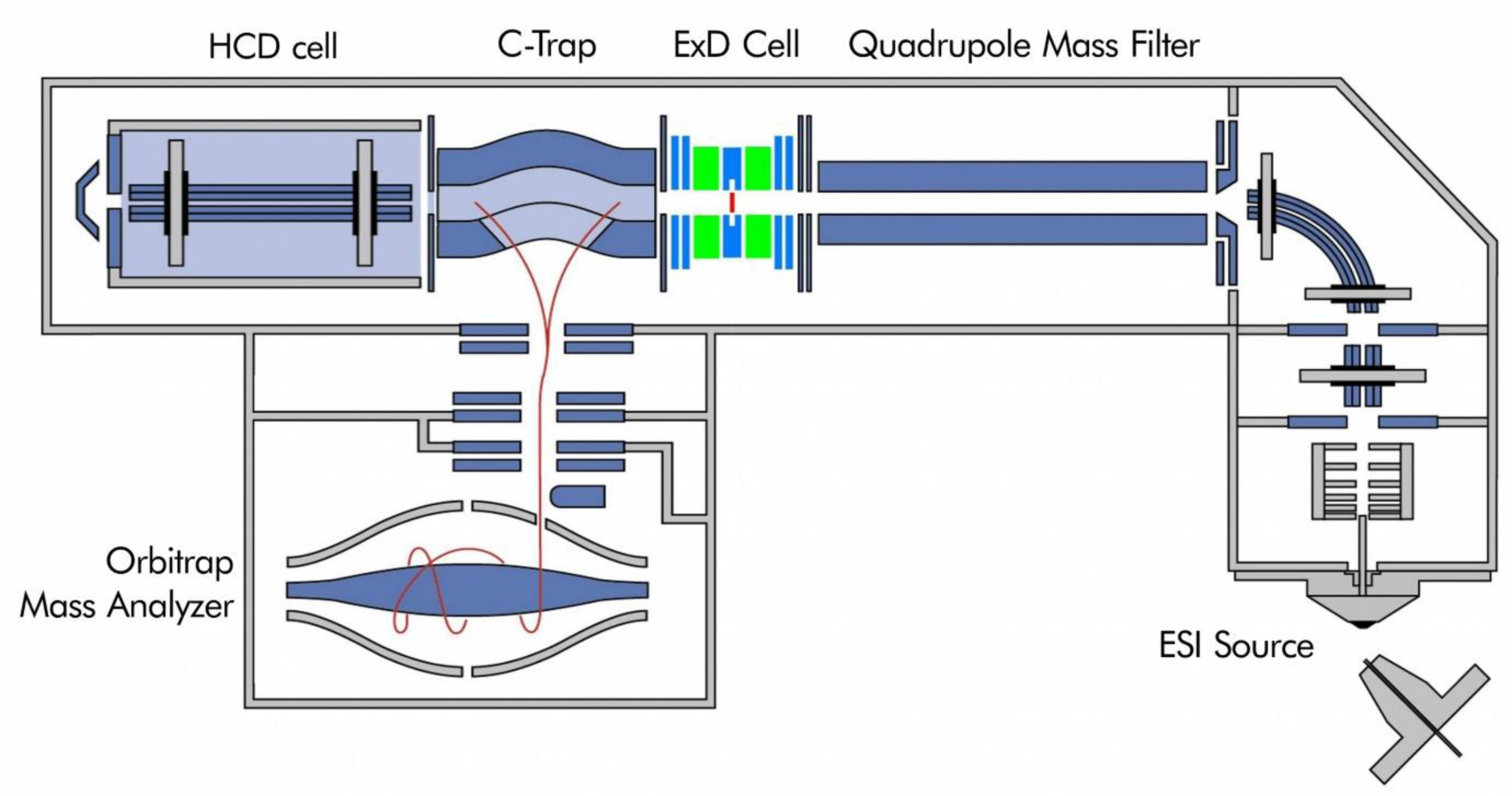
Schematic of the Q Exactive UHMR MS with the installed ExD TQ-160 cell. The ExD cell replaces the transfer multipole in this instrument.

**Suppl. Fig. 5.**
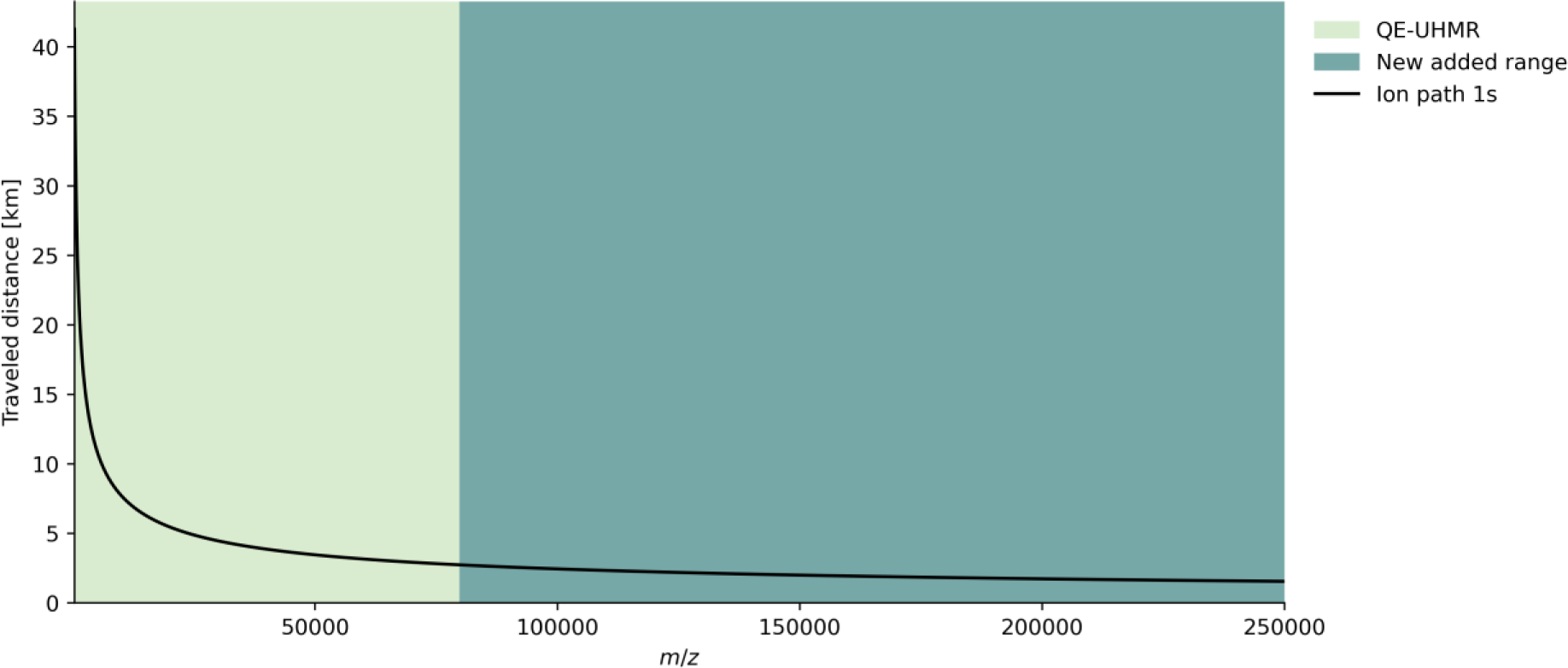
Distance travelled vs. m/z for ions during a 1 second transient. Higher m/z ions, having lower frequencies, travel reduced distances in the Orbitrap analyser at lower velocities compared to lower m/z ions over the same transient time, resulting in fewer and less energetic collisions. The plot background is coloured to illustrate the standard m/z range of the Q Exactive UHMR MS and the further range newly added in this work.

